# Type-specific EV-D68 real-time RT-PCR assay for the detection of all extant enterovirus D68 strains

**DOI:** 10.1101/2022.10.06.511205

**Authors:** Terry Fei Fan Ng, W. Allan Nix, Shannon L. Rogers, Brian Emery, Shur-Wern Chern, Kiantra Butler, M. Steven Oberste

## Abstract

Enterovirus D68 (EV-D68) has caused recurring respiratory disease outbreaks in the United States since 2014. The dominant circulating EV-D68 strain has evolved from clade B1 to the more recent B2 and B3 clades. As recurrent outbreaks and continued virus evolution are expected for EV-D68, a robust real-time PCR assay that detects known strains as well as potential emerging strains is critical for national surveillance and clinical diagnostics. We describe a type-specific EV-D68 real-time RT-PCR (rRT-PCR) assay termed CDC2022, which targets sequences encoding conserved amino acid regions of all extant EV-D68 strains. We targeted three motifs conserved among all strains in the last 60 years. The assay achieved 100% (270/270) sensitivity and 100% (344/344) specificity when tested with a collection of 613 respiratory specimens, compared to the gold-standard EV semi-nested VP1 PCR and sequencing assay (snPCR/Seq). CDC2022 gave negative results with 289/289 non-target viruses, including 104 EV A-D isolates, 165 rhinovirus (RV) isolates or clinical specimens, and 14 other common respiratory viruses. The assay can detect as few as 0.28 CCID_50_ per reaction. An *in silico* “phylo-primer-mismatch” analysis was performed to visualize primer/probe mismatches and to compare CDC2022 with other EV-D68 rRT-PCR assays, including the previous CDC assay (CDC2015) developed in 2014 for clade B1 strains. It showed that CDC2022 has the fewest primer/probe mismatches among all assays analyzed and is suitable for all clades. We additionally tested 11 EV-D68-positive clinical specimens from 2022 that were confirmed by snPCR/Seq, and all were detected. CDC2022 assay could provide a critical tool for molecular surveillance of EV-D68.

## Introduction

Enterovirus D68 (EV-D68; genus *Enterovirus*, family *Picornaviridae*) can cause severe respiratory illness, with clinical manifestations that include bronchiolitis, wheezing, and pneumonia, especially in children (1). EV-D68 has also been associated with acute flaccid myelitis (2–5). EV-D68 had been relatively rare from its discovery in 1962 until the mid-2000s (6) but more recently, it has been associated with clusters and outbreaks of severe respiratory disease worldwide (2, 7, 8). In 2014, EV-D68 caused a nationwide outbreak of severe respiratory tract illness in the United States, with over 1100 laboratory-confirmed cases in 49 states and the District of Columbia. Successive outbreaks have occurred in the US and many other countries in 2016 and 2018, suggesting a biannual pattern. A low level of circulation has been observed since 2019 (7), likely due to non-pharmaceutical interventions instituted to prevent the spread of SARS-CoV-2 (9).

Enteroviruses (EVs) other than EV-D68 do not usually cause respiratory diseases. At CDC, pan-EV real-time reverse-transcription polymerase chain reaction (rRT-PCR)(10) and EV VP1 semi-nested PCR and sequencing assays (snPCR/seq) (11) are the primary approaches for enterovirus diagnostics. While the gold-standard snPCR/seq method allows sensitive detection of all enteroviruses and many rhinoviruses, as well as identification of virus type, it is labor-intensive and difficult to scale to large numbers of specimens. To facilitate more rapid laboratory identification of EV-D68 cases during the 2014 outbreak investigation, an EV-D68-specific real-time PCR (rRT-PCR) assay was developed (https://www.fda.gov/media/120427/). It targets the dominant strain that circulated in 2014, which is now defined as clade B1. Clade B1 continued to evolve, so by the time of the 2016 and 2018 outbreaks, the major circulating lineages were clades B2 and B3, but a small number of clade D cases have also been reported. Clade A has not been detected post-2014. As the sequences of the predominant circulating strains continue to evolve, the performance of CDC2015 has decreased (12, 13), likely due to sequence divergence compared to the original 2014 clade B1 outbreak strain. Other rRT-PCR assays, from Washington University (WU) (14) and from Niigata University (NU) (12) have been described for detection of the newer clades.

With the recurrence of EV-D68 circulation and continued evolution of the virus, a versatile EV-D68 rRT-PCR assay is needed to detect not only all known clades, but also future, potentially divergent strains. Here, we describe a *pan*-EV-D68 rRT-PCR assay (called CDC2022) that detects all known EV-D68 clades. In addition to validation of the CDC2022 assay, we present *in silico* analyses of the CDC2015, CDC2022, WU, and NU assays to predict clade-specific characteristics of these commonly used EV-D68 rRT-PCR assays.

## Materials and Methods

### Primer and probe design

To design the pan-EV-D68 primers (CDC2022), EV-D68 VP1 protein sequences were analyzed to identify regions of amino acid conservation (enterovirus VP1 contains immunodominant neutralization epitopes and its sequence correlates with serotype (15). Representative sequences from EV-D68 clades A, B, C, and D were obtained from GenBank and sequences were manipulated using Geneious (versions including R9 and 2022.1.1). Only VP1-containing sequences with over 900 bp length were selected for further analysis. To reduce duplicative data, VP1 sequences with >99% nucleotide identity (NI) were excluded. VP1 nucleotide sequences were translated, and the amino acid sequences were aligned using MAFFT (16). Sequence logo and sliding window analysis with a window size of 30 amino acids were performed using Geneious to identify conserved amino acid motifs. As real-time PCR requires primers and a probe in close proximity to one another, selection of primer locations prioritized areas that are both conserved and close enough to one another to accommodate the constraints of a real-time PCR assay. Primer and probe sequences were manually designed on targeted conserved VP1 motifs using the nucleotide sequence alignment (Fig. 1), and primer properties including potential for primer-dimer formation were checked using Oligo Analyzer (IDTDNA).

**FIG 1.**
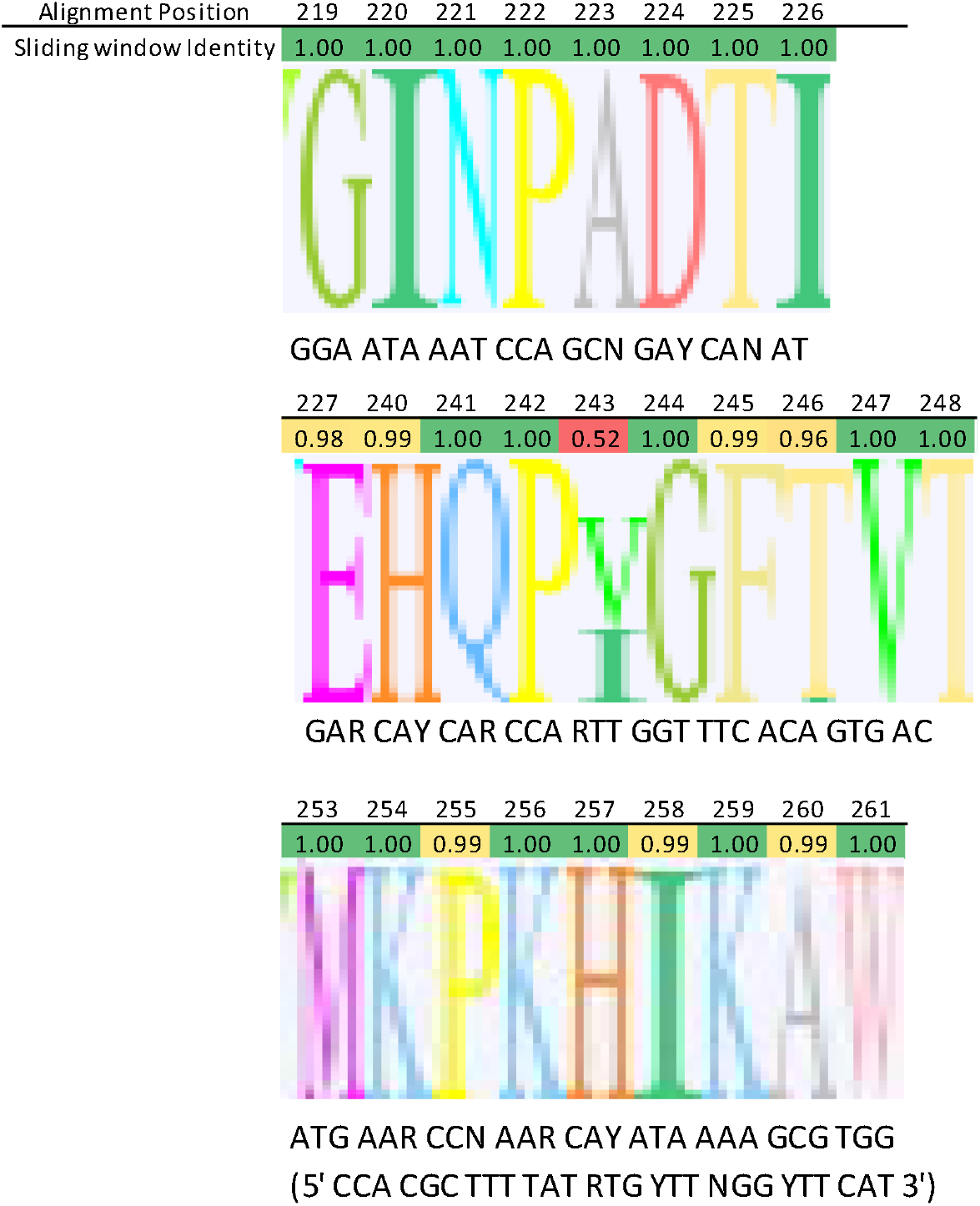
Conserved motifs targeted for CDC2022 pan-EV-D68 rRT-PCR.

### Real-time RT-PCR conditions

rRT-PCR for CDC2022 was performed using the qScript™ XLT One-Step RT-qPCR ToughMix® (Quanta Biosciences; catalog #95132-500 or 95132-100). The reaction components and thermocycling conditions are provided in detail in Tables S1 and S2. Briefly, the 20 μl reaction mixture contained 10 μl of ToughMix enzyme reagent, 0.5 μl of 20 μM AN993 sense primer, 0.5 μl of 20 μM AN995 antisense primer, and 1 μl of 10 μM AN992 probe (0.5 μM final concentration of each primer and the probe), 3 μl of sterile water, and 5 μl of input RNA. Thermocycling and detection were performed using a 7500 Fast Real-Time PCR System (Applied Biosystems) according to the manufacturer’s instructions.

### RNA extraction

Virus isolates and respiratory specimens were extracted using the QIAgen Viral RNA Mini-Kit (QIAgen, Inc., Valencia, CA). Input sample volume was 140 μl, and RNA was eluted in 60 μl sterile nuclease-free water. Extracted nucleic acids were stored at −20°C until testing.

### Virus strains

EV-D68 isolates used in this study included Fermon prototype (GenBank accession number AY426531), US/MO/14-18947 (KM851225), US/MO/14-18949 (KM851227), US/IL/14-18952 (KM851230) and US/KY/14-18953 (KM851231)(17). EV-D68 isolates were grown in RD cells in 25 cm^2^ flasks and harvested at 60% to 70% cytopathic effect. The infected RD cells were frozen and-thawed twice, and the supernatants were collected and clarified by centrifugation at 10,000 x *g* for 10 minutes, before aliquoting and freezing at −70°C. An aliquot of the virus was thawed and titrated on RD cell monolayers to determine end point titer, expressed in units of 50% cell culture infectious dose (CCID_50_).

### Analytical sensitivity testing

To facilitate sensitivity evaluation, the Fermon prototype strain and representative EV-D68 isolates from clades B1, B2, B3, and D were spiked into minimum essential medium (MEM). Isolates were titrated before spiking. The RNA was then extracted and serially diluted to the range of 10^−5^ to 10^−9^ For each strain, each dilution was tested three times using the CDC2022 assay. Similar sensitivity evaluation was performed with the CDC2015 assay (Table S3) with a smaller isolate collection. The CDC2022 assay sensitivity was further evaluated by spiking two representative isolates into EV-negative clinical nasopharyngeal/oropharyngeal swab matrix, with 20 technical replicates. Titrated clade B1 isolate US/MO/14-18949, and clade B2 isolate US/18-23087 were first serially diluted to the range of 10^−6^ to 10^−8^, then dilutions were independently extracted 20 times and tested using the CDC2022 assay.

### Analytical specificity testing

The analytical specificity of CDC2022 was evaluated against viruses from *Enterovirus* species A-D and *Rhinovirus* species A-C, and thirteen other common respiratory viruses (summarized in Table 4, enumerated in Tables S4-S7). The EV A-D samples are isolates collected in the CDC enterovirus reference laboratory, including 22 EV-A, 56 EV-B, 17 EV-C, and 6 EV-D isolates. The closest sequence relatives to EV-D68 are other EV-D types, and currently four other types were known. Three of the four non-EV-D68 EV-D types were tested (EV-D70, EV-D94, and EV-D111); EV-D120, isolated from gorillas and a chimpanzee, in Cameroon and the Democratic Republic of Congo, respectively, was not available for testing. To date, EV-D120 has not been detected in humans. In addition to the available “nearest neighbors” in EV-D, other common respiratory viruses were also tested. These included 95 rhinovirus (RV) isolates from species A and B (RV-A, RV-B), representing the classically defined serotypes, as well as adenovirus C1, coronaviruses (229E, OC43, and MERS-CoV), human metapneumovirus, influenza virus (A H1N1 and A H3N2, and B), parainfluenza viruses 1-3 and 4a, respiratory syncytial virus, measles virus, and mumps virus. Rhinovirus species C (RV-C) (genus Enterovirus) have not been successfully grown in conventional cell cultures. RNA was extracted from 69 respiratory tract clinical specimens (nasopharyngeal/oropharyngeal swabs in viral transport medium) in which RV-C were identified by direct sequencing during 2014 and used to test CDC2022 assay specificity.

### Phylo-primer-mismatch analysis

Similar to the approach used for primer design, representative EV-D68 sequences were obtained from GenBank, and were manipulated using Geneious (versions including R9 and 2022.1.1). VP1 sequences over 900 bp length were selected for analysis and VP1 sequences with >99% nucleotide identity (NI) were excluded to reduce duplication. The 359 unique VP1 nucleotide sequences were aligned using MAFFT (16). For each assay, primer and probe sequences were mapped against the VP1 alignment to analyze the number of mismatches per strain. A neighbor-joining tree using the Jukes-Cantor substitution model was generated using Geneious. The mismatches were tabulated for visualization and overlayed with the phylogenetic tree.

### Testing of clinical specimens

A total of 614 respiratory specimens collected in 2012-2019 were tested in parallel using snPCR (gold standard) (11) and CDC2022 (Table S9). The partial VP1 sequences by snPCR were generated using the snPCR/Seq method (11), and typed using CDC PiType (https://pitype.cdc.gov/). Eleven 11 specimens collected in 2022 and all ultimately identified as EV-D68 (Table S9) were also tested by the same methods.

## Results

### Primer and probe design

A comprehensive collection of EV-D68 VP1 clades A-D VP1 sequences were analyzed for conserved motifs suitable for a pan-EV-D68 rRT-PCR assay. Three neighboring conserved motifs, GINPADTI (sense primer), MKPKHIKAW (antisense primer), and EHQP(V/I)GFTVT (probe) were identified (Fig. 1). Each of these motifs is conserved for >99% of all available EV-D68 VP1 sequences. Sequence logo and sliding window analysis confirmed the same degree of sequence conservation (Fig. 1). Nucleotide degeneracy is incorporated in the 3’ half of the primers to account for all codons of the target protein motifs. The primer and probe sequences are described in Table 1.

**Table 1.**
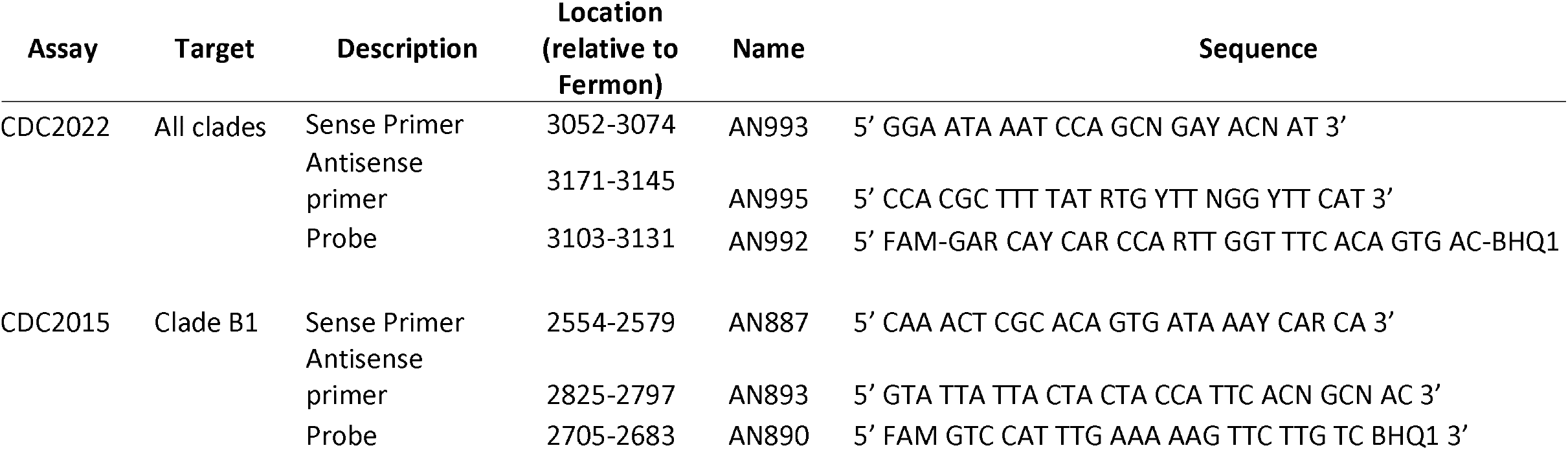
EV-D68 rRT-PCR CDC2022 and CDC2015 assays primers and probes. Positions were relative to EV-D68 prototype strain (Fermon; GenBank accession AY426531). Nucleotide degeneracy was noted according to IUPAC ambiguity codes.

### Phylo-primer-mismatch analysis to assess primer mismatches to predict assay performance

The primer and probe sequences for the CDC2015 and CDC2022 assays and other published EV-D68 assays (12, 14) were aligned with 359 unique, representative EV-D68 VP1 sequences (sequences with >99% nucleotide identities were deduplicated). We developed a “phylo-primer-mismatch graph,” depicting phylogenetic relationships among the reference sequences and degree of primer or probe mismatch, to visualize the number of sequence mismatches for each assay (Fig. 2). Sense primer, antisense primer, and probe were evaluated for each assay.

**FIG 2.**
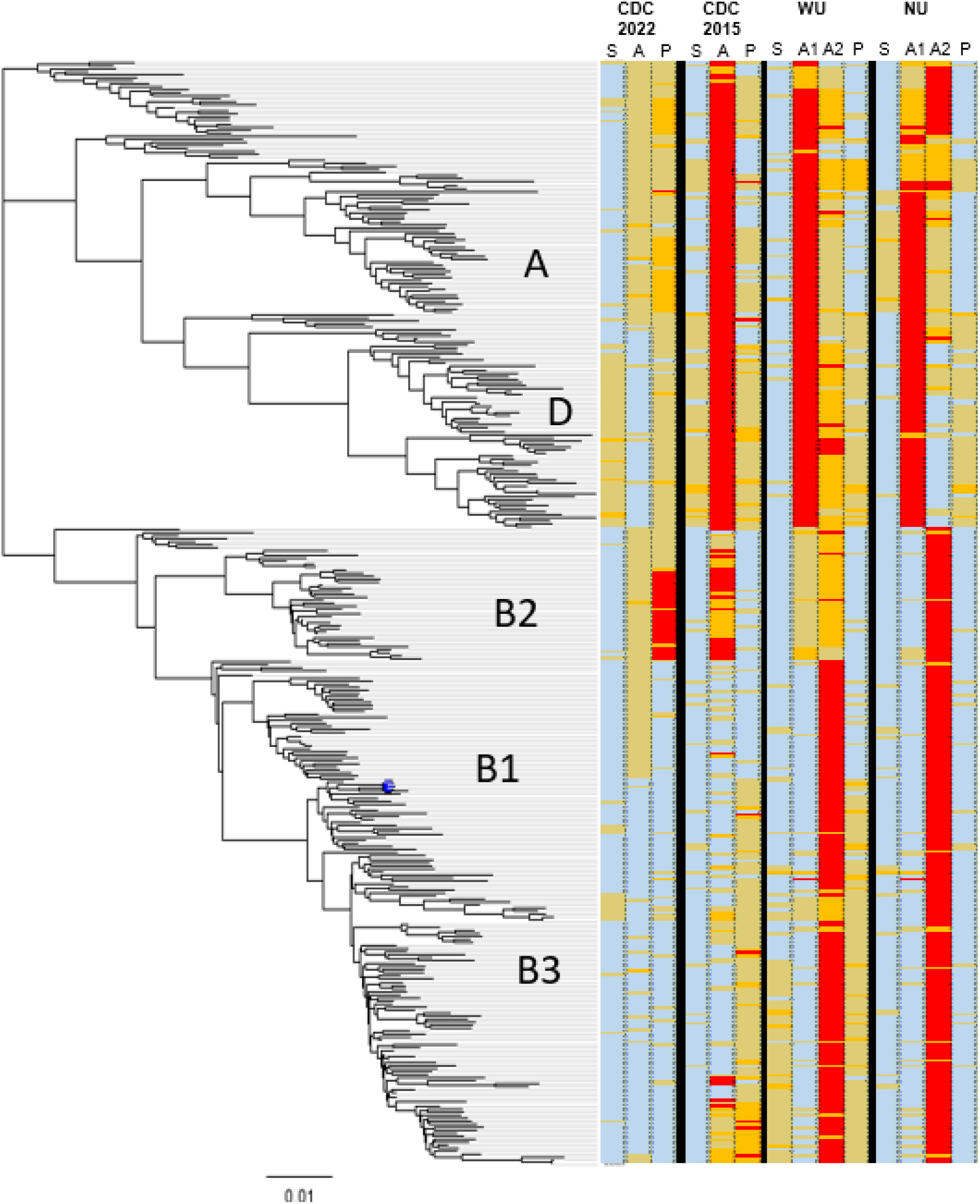
Phylo-primer-mismatch graph for visualization of primer/probe mismatches against the EV-D68 phylogeny. Alignment of 359 representative deduplicated VP1 nucleotides were used to construct a neighbor-joining tree. Primer/probe mismatches to each sequence were tabulated and overlayed with the tree. Sense primer, antisense primer, and probe were evaluated in this order for each assay. WU and NU assays have two antisense primers. S, sense primer; A, antisense primer; A1, antisense-1 primer; A2, antisense-2 primer; P, probe. Blue, 0 mismatch; Yellow, 1 mismatch; Orange, 2 mismatches; Red, 3 or more mismatches.

The CDC2022 sense and antisense primers and probe contain zero mismatches for most clade B strains, except for 27 clade B2 sequences that had 3-4 probe mismatches. They have ≤2 mismatches with clade A and D strains. In comparison, the CDC2015 assay has zero mismatches to most of the targeted clade B1 (range 0-3) but it contains 3-12 mismatches to sequences from clades A and D, and to historical strains from 1962 to the 2000s, including Fermon, as well as some clade B2 strains.

The phylo-primer-mismatch graph approach can visualize the primer-target relationships of existing assays, to better understand their design. Both WU and NU assays contain two antisense primers (Fig. 2). The antisense-1 primer targets the clade B strains, while antisense-2 primer targets clades A and D and the pre-2014 strains such as Fermon. Because the dual antisense primers allow either primer to bind to the target, the lowest mismatch of the two antisense primers should be considered when comparing primer/probe binding. For the WU assay, the antisense-1 primer targeting clade B contains zero mismatches to most clade B sequences (range 0-3). However, the antisense-2 targeting clade A and D contains two mismatches (range 0-3). The WU sense primer and probe contain majority of 0 mismatch with clades B1 and B2 but had more mismatches (majority of 1-2 mismatches) with sequences in clade B3. The NU assay sense primer, antisense-1 primer, and probe have zero mismatches with most clade B sequences (range 0-3). The probe has one mismatch with most clade D sequences. The antisense-2 primer has more mismatches (1–4) with clade A and pre-2014 strains. Considering matched sequences with their primer/probe, the highest number of mismatches per virus sequence for CDC2022, CDC2014, WU, and NU are 4, 12, 7, and 6, respectively.

### Analytical sensitivity and specificity

The assay analytical sensitivity was evaluated by spiking the Fermon strain and representatives of EV-D68 clades B1, B2, B3, and D (Table 2) into MEM, followed by serial dilution and testing in the CDC2015 and CDC2022 assays. In MEM, the CDC2022 assay consistently detected as few as 10 CCID_50_/ml (0.28 CCID_50_ per reaction) of EV-D68 clades B1, B2, B3, D, and the Fermon strain (Table 2). The CDC2015 assay showed similar performance with clade B isolates (Table S3), but CDC2015 is at least 10 times less sensitive than CDC2022 for clade D and Fermon, likely due to the large number of mismatches identified in the phylo-primer-mismatch analysis.

**Table 2.** Analytic sensitivity of the CDC2022 EV-D68 rRT-PCR assays. Extracted isolate RNA was serially diluted and tested three times per strain. Virus titer (CCID_50_/ml) after dilution factor is shown in the parenthesis.

| Assay | RNA<br>Dilution | Fermon<br>Prototype<br>(historical) | US/IL/14-18952<br>(clade B1) | US/MO/14-18949<br>(clade B2) | US/2018-23087<br>(clade B3) | US/KY/14-18953<br>(clade D) |
| --- | --- | --- | --- | --- | --- | --- |
| CDC2022 | undiluted<br>stock CCID50 | 10<sup>5.9</sup> | 10<sup>8.0</sup> | 10<sup>7.8</sup> | 10<sup>7.1</sup> | 10<sup>7.1</sup> |
|  | 10-5 | 3/3 (8) | 3/3 (1000) | 3/3 (630) | 3/3 (125) | 3/3 (125) |
|  | 10-6 | **3/3 (0.8)** | 3/3 (100) | 3/3 (63) | **3/3 (12.5)** | 3/3 (12.5) |
|  | 10-7 | 0/3 (0.08) | **3/3 (10)** | **3/3 (6)** | 3/1 (1.25) | **3/3 (1.25)** |
|  | 10-8 | 0/3 (0.008) | 1/3 (1) | 2/3 (0.6) | 0/3 (0.125) | 1/3 (0.125) |
|  | 10-9 | 0/3 (0.008) | 0/3 (0.1) | 0/3 (0.06) | 0/3 (0.0125) | 0/3 (0.0125) |

The CDC2022 assay was tested using virus-spiked enterovirus-negative respiratory specimen matrix (nasopharyngeal/oropharyngeal swab in viral transport medium). The CDC2022 assay detected down to 63 CCID_50_/ml (0.735 CCID_50_ per reaction) for clade B1 strain 14-18949 (100% of 20 replicates) and 13 CCID_50_/ml (0.146 CCID_50_ per reaction) for clade B3 strain 18-23087 (100% of 20 replicates) (Table 3).

**Table 3.** Reproducibility and limit of detection (LOD) of the EV-D68 rRT-PCR CDC2022 assay determined using spiked clinical matrix (NP/OP). The pooled matrix spiked with isolates was serially diluted. Each dilution was extracted 20 times and tested.

**EVD-68 virus isolate US/MO/14-18949**
| **Dilution** | **CCID<sub>50</sub>/ml isolate** | **CCID50 per rRT-PCR** | **Call Rate** | **Call Rate %** | **Average Ct** | **Ct SD** |
| --- | --- | --- | --- | --- | --- | --- |
| 10<sup>-6</sup> | 63 | 0.7350 | 20/20 | 100% | 32.8 | 0.69 |
| 10<sup>-7</sup> | 6.3 | 0.0735 | 16/20 | 80% | 36.4 | 2.1 |
| 10<sup>-8</sup> | 0.63 | 0.0074 | 1/20 | 5% | NA | NA |

**EVD-68 virus isolate US/18-23087**
| **Dilution** | **CCID<sub>50</sub>/ml isolate** | **CCID50 per rRT-PCR** | **Call Rate** | **Call Rate %** | **Average Ct** | **Ct SD** |
| --- | --- | --- | --- | --- | --- | --- |
| 10<sup>-6</sup> | 12.5 | 0.1458 | 20/20 | 100% | 32.2 | 0.55 |
| 10<sup>-7</sup> | 1.25 | 0.0146 | 14/20 | 70% | 37.1 | 1.83 |
| 10<sup>-8</sup> | 0.125 | 0.0015 | 4/20 | 20% | 39 | 2.14 |

The analytical specificity of the pan-EV-D68 rRT-PCR (CDC2022) was evaluated by testing against a broad panel of viruses other than EV-D68, including *Enterovirus* species A-D (non-EV-D68 in EV-D) (Table S4), *Rhinovirus* species A-C (Tables S5 and S6), and 14 other common respiratory viruses (Table S7). All 289 non-EV-D68 viruses were negative using the CDC2022 assay, for an analytical specificity of 100% (summarized in Table 4).

**Table 4.** Analytic specificity of EV-D68 rRT-PCR CDC2022 assay with enteroviruses, rhinoviruses, other respiratory viruses. The viruses tested are described in detail in Tables S1-S4.

| Cell Culture Isolates or Respiratory Clinical Specimens | Summary |  |
| --- | --- | --- |
|  | Negative/Total | Specificity (%) |
| EV-D species | 6/6 | 100 |
| RV-A, -B and -C species | 165/165 | 100 |
| EV-A, -B and -C species | 104/104 | 100 |
| Other Common Respiratory Viruses | 14/14 | 100 |
| **Overall Analytical Specificity** | **289/289** | **100** |

### CDC pan-EV-D68 rRT-PCR analytical sensitivity with clinical specimens

To evaluate the sensitivity of the CDC2022 assay in clinical specimens, we tested 614 respiratory specimens using the CDC2022 pan-EV-D68 rRT-PCR and the EV VP1 sequencing assay (snPCR/Seq). All 270 specimens that were EV-D68-positive by snPCR/Seq were detected by the CDC2022 assay, attaining 100% sensitivity (Fig. 3). The 344 snPCR/Seq EV-D68-negative samples (a combination of other enteroviruses and samples that were negative for all enteroviruses) were also negative in the CDC2022 assay (Suppl Table S9). The CDC2022 assay successfully detected EV-D68 in 11 clinical specimens from 2022 in which EV-D68 was confirmed by snPCR and sequencing as clade B3, the predominant clade that circulated during 2022 (Ng et al., unpublished data).

**FIG 3.**
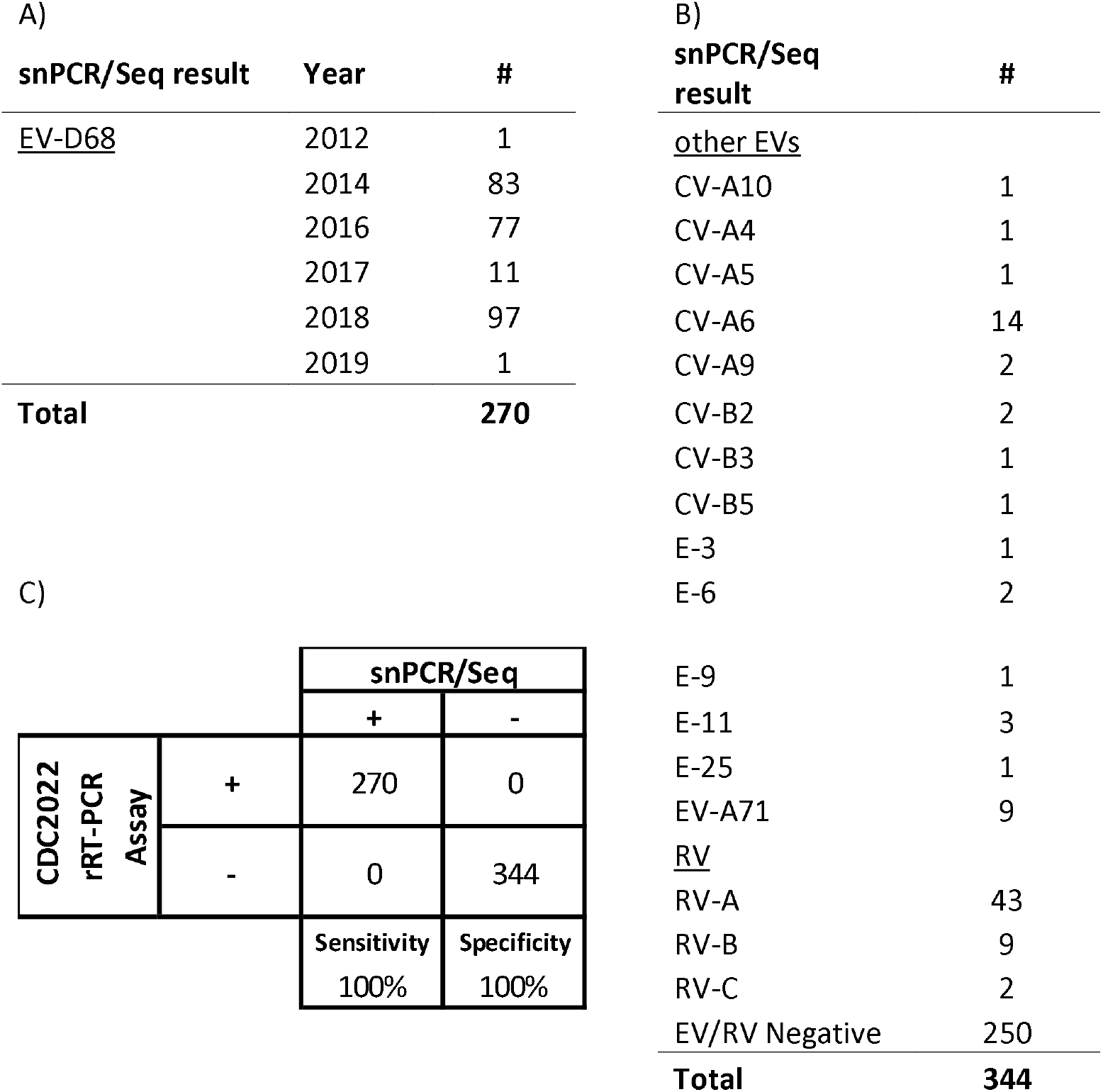
Clinical sensitivity of CDC-2022 rRT-PCR compared against the gold-standard snPCR/Seq. A) Breakdown of the EV-D68 positive specimens tested by snPCR/Seq. B) Breakdown of all other specimens tested and their snPCR/Seq result. C) Confusion matrix of the rRT-PCR compared to snPCR/Seq showing the assay’s sensitivity and specificity in clinical specimens described in A) and B).

## Discussion

Enterovirus infections are most frequently diagnosed by real-time RT-PCR targeting the highly conserved 5’-non-translated region of the viral genome (10). In most cases, identification of enterovirus type is not necessary for clinical management of enterovirus disease and a result of “EV-positive” is a sufficient result, if only to help rule out other etiologies. Unfortunately, pan-EV assays tend to cross-react with rhinoviruses (and vice versa), making it difficult to differentiate between rhinovirus and enterovirus infection in respiratory disease (18). As a result, the available FDA-approved molecular diagnostic tests for testing of respiratory specimens usually report the result as “RV/EV positive.” However, enterovirus typing can be helpful in identifying disease clusters or outbreaks, or when patients are being cohorted for infection control. In such cases, the method of choice is PCR-amplification of a portion of the region that encodes the VP1 capsid protein, a region whose sequence correlates with antigenic type, followed by sequencing (11). While current VP1 PCR/sequencing methods are sensitive and can yield molecular epidemiologic data in addition to virus type, these methods are relatively labor-intensive and do not scale well to large numbers of specimens. Such assays are also not widely available in state or local public health laboratories.

Type-specific real-time RT-PCR assays have been developed for several enteroviruses, including polioviruses (19), enterovirus A71 (20, 21), and coxsackievirus A16 (21). The major application of such assays is in disease surveillance to rapidly identify a specific virus of interest, such as in acute flaccid paralysis surveillance in support of global polio eradication (19) or during an outbreak in which a specific virus type is suspected or has been identified as the major etiologic agent (21). The major advantages of real-time PCR assays over sequencing are speed and scalability, since an entire 96-well PCR plate can be quickly set up and run in only a few hours. This can be essential in an outbreak situation, such as the 2014 US EV-D68 outbreak, where the number of specimens can rise quickly, overwhelming the diagnostic laboratory. Since modern virus diagnostic laboratories are likely to already have real-time PCR instrumentation and trained staff for other virus diagnostic assays, the real-time PCR format makes it easy to rapidly deploy a new assay in diagnostic and public health laboratories.

The CDC2015 assay was developed at the peak of the 2014 EV-D68 respiratory disease outbreak in the US to facilitate more rapid differentiation of EV-D68 outbreak cases from the background of sporadic disease due to rhinoviruses and other enteroviruses (https://www.fda.gov/media/120427/). The CDC2015 assay sense and antisense primers had four-fold and 16-fold sequence degeneracy, respectively, to account for nucleotide variation in the outbreak strains and related historical sequences; the probe sequence is non-degenerate. For that assay, we made a design decision toward absolute sensitivity at the expense of breadth of reactivity, so the CDC2015 assay could substitute for the sensitive gold standard assay, snPCR/Seq. As a result, the assay can detect EV-D68 sequences in clade B, but not the single outbreak virus that was in an outlier cluster (17), now known as Clade D. We considered this to be a reasonable compromise at the time, given that >99% of EV-D68 sequences from the 2014 outbreak were in clade B and were detected by the CDC2015 assay.

Since 2016, the major circulating EV-D68 strains have evolved to become clades B2 and B3, and several newer rRT-PCR assays were developed to detect the emerging strains (12, 14), yet some of the assays were shown to be less sensitive for certain clades (12). With the recurrence of EV-D68 outbreaks in recent years, we aimed to design a pan-EV-D68 rRT-PCR assay that detects all clades. By targeting conserved sites in the VP1 region, the assay should remain robust for detection of future EV-D68 sequence divergence. The sites encoding the GINPADTI amino acid motif (sense primer), the MKPKHIKAW motif (antisense primer), and the EHQP(V/I)GFTVT motif (probe) have been conserved among >99% of EV-D68 strains for the past 60 years, including the prototype strain, Fermon, and the most recent viruses in clade B3. The phylo-primer-mismatch analysis showed that the CDC2022 assay has 3-4 mismatches with several sequences from clade B2, but upon inspection, most of the mismatches are in the 5’ half of the probe and therefore have a lesser effect on probe binding. We believe the CDC2022 assay will provide a critical tool for molecular surveillance of EV-D68 as the virus continues to evolve, though ongoing strain surveillance will be required to ensure that is the case.

We developed a “phylo-primer-mismatch analysis” approach to visualize predicted primer and probe performance against a group of EV-D68 strains to assess the EV-D68 assays, but the approach could be applied to other pathogens as well. Although mismatches could be reported in table format, the graphic overview provides useful insight into whether the mismatch(es) will affect primer/probe performance with specific virus clades. Through this analysis, we demonstrated the CDC2022 assay’s superiority over the CDC2015 assay in breadth of target range, in agreement with the assays’ performance in clinical testing. Primer mismatch analysis should not replace actual “wet-lab” comparison of assay performance, but it provides *in silico* evidence to guide assay design and to better understand observed assay performance. As currently implemented, the phylo-primer-mismatch analysis is largely a manual process, but a bioinformatic script that automates the analysis could be developed. Beyond enteroviruses, such a tool could help evaluate novel assay designs against an emergent strain or clade of a pathogen, to ensure the primers and probe(s) are suitable for detecting all intended targets and account for sequence divergence. It can also be used to evaluate existing primers and probes against an ever-evolving virus.

This paper describes evaluation of CDC2022 rRT-PCR assay performance with a large specimen set for assessment of clinical sensitivity and specificity (n=614) and analytical specificity (n=289), providing validation of the assay’s robustness. *In silico* analysis showed CDC2022 has the fewest primer/probe mismatches among all assays evaluated. The NU assay also has very few primer/probe mismatches with the currently circulating B and D clades, which was supported by good assay performance (12), suggesting it is likely to perform well for currently circulating strains. However, unlike CDC2022, the NU and WU primers are not designed to account for target sequence evolution, and therefore they may fail to detect future EV-D68 strains. The CDC2022 assay targeted amino acid motifs that have been conserved for decades, making the assay resistant to future sequence divergence.

While rRT-PCR is an efficient way to diagnose EV-D68 in clinical cases, it does have limitations. The amplicon is too short for sequencing, and therefore not ideal for molecular epidemiology or clade determination. Whole-genome amplification techniques (22), which may be more easily automatable than snPCR/seq, can be used as a reflex test for the rRT-PCR-positive specimens to generate genomes for molecular analysis. The whole genome sequences, in turn, can help to identify potential primer or probe mismatches and allow further improvement of the rRT-PCR assay.

## Acknowledgements

“The findings and conclusions in this report are those of the author(s) and do not necessarily represent the views of CDC and other contributing agencies.” The use of trade names is for identification purposes only and does not constitute an endorsement by the Centers for Disease Control and Prevention or US Government. We thank Kaija Maher for excellent technical assistance in evaluating the assays.

