## Supplemental File for "Type-specific EV-D68 real-time RT-PCR assay for the detection of all extant enterovirus D68 strains" (from another version of this preprint: version 1, 2022-10-06)

### Supplemental materials

**Table S1.** rRT-PCR reaction conditions for Applied Biosystems 7500 Fast.

| Component | Single reaction |
|---|---|
| **2X ToughMix** enzyme reagent | 10.0 µL |
| AN993 primer (20 µM) | 0.5 µL |
| AN995 primer (20 µM) | 0.5 µL |
| AN992 probe (10 µM) | 1.0 µL |
| Sterile nuclease-free water | 3.0 µL |
| Extracted nucleic acid | 5.0 µL |
| Total volume | 20.0 µL |

**Table S2.** Thermocycling profile

| Step | Cycles | Temperature | Time |
|---|---|---|---|
| Reverse transcription | 1 | 50°C | 30 min |
| Taq inhibitor inactivation | 1 | 95°C | 1 min |
| Polymerase chain reaction | 45 | 95°C<br>52°C | 15 sec<br>50 sec |

**Table S3**. Analytic sensitivity of the EV-D68 rRT-PCR CDC2022 and CDC2015 assays.

| **Assay** | **RNA Dilution** | **Fermon Prototype (historical)** | **US/IL/14-18952 (clade B1)** | **US/MO/14-18949 (clade B2)** | **US/2018-23087 (clade B3)** | **US/KY/14-18953 (clade D)** |
|---|---|---|---|---|---|---|
| **CDC2022** | | | | | | |
| | **undiluted stock CCID50** | **$10^{5.9}$** | **$10^{8.0}$** | **$10^{7.8}$** | **$10^{7.1}$** | **$10^{7.1}$** |
| | **10-5** | 3/3 (8) | 3/3 (1000) | 3/3 (630) | 3/3 (125) | 3/3 (125) |
| | **10-6** | **3/3 (0.8)** | 3/3 (100) | 3/3 (63) | **3/3 (12.5)** | 3/3 (12.5) |
| | **10-7** | 0/3 (0.08) | **3/3 (10)** | **3/3 (6)** | 3/1 (1.25) | **3/3 (1.25)** |
| | **10-8** | 0/3 (0.008) | 1/3 (1) | 2/3 (0.6) | 0/3 (0.125) | 1/3 (0.125) |
| | **10-9** | 0/3 (0.008) | 0/3 (0.1) | 0/3 (0.06) | 0/3 (0.0125) | 0/3 (0.0125) |
| **CDC2015** | | | | | | |
| | **10-5** | 0/3 (8) | 3/3 (1000) | 3/3 (630) | not tested | **3/3 (125)** |
| | **10-6** | 0/3 (0.8) | 3/3 (100) | 3/3 (63) | not tested | 0/3 (12.5) |
| | **10-7** | 0/3 (0.08) | **3/3 (10)** | 3/3 (6) | not tested | 0/3 (1.25) |
| | **10-8** | 0/3 (0.008) | 2/3 (1) | **3/3 (0.6)** | not tested | 0/3 (0.125) |
| | **10-9** | 0/3 (0.008) | 0/3 (0.1) | 1/3 (0.06) | not tested | 0/3 (0.0125) |

### CDC2022 assay specificity

**Enterovirus species A, B and C.** Most EV types are not thought of as frank respiratory pathogens; however, most EV species A, B, and C do replicate to some extent in the nasopharynx and some types other than EV-D68 can cause acute respiratory disease, e.g., CV-A21 and EV-A71. Because of this, we investigated all the human (and non-human primate) EV species A-C for cross reactivity (**Table S1**). RNA was extracted from cell culture supernatants and tested undiluted. The Pan-EV-D68 rRT-PCR did not amplify any viruses from EV-A, EV-B, or EV-C.

**Enterovirus Species D.** To evaluate the analytical specificity of the Pan-EV-D68 rRT-PCR, we tested a panel of enterovirus species D (EV-D; five known types including EV-D68) cell culture isolates. Respiratory clinical specimens positive for EV-D70, EV-D94, EV-D111 were not available for testing, therefore RNA from these viruses was extracted from frozen undiluted cell culture supernatants and tested. The Pan-EV-D68 rRT-PCR did not amplify other *Enterovirus D* viruses (**Table S1**).

**Table S4 Pan-EV-D68 rRT-PCR analytical specificity confirmed using undiluted RNA extracted from enterovirus cell culture supernatants**

| Species | Enterovirus Types | Pan-EV-D68 rRT-PCR Result |
|---|---|---|
| **EV-A** | Coxsackievirus A2 (CV-A2), CV-A3, CV-A4, CV-A5, CV-A6, CV-A7, CV-A8, CV-A10, CV-A12, CV-A14, CV-A16, Enterovirus A71 (EV-A71), EV-A76, EV-A89, EV-A90, EV-A91, EV-A92, EV-A114, EV-A122, EV-A123, EV-A124, EV-A125 | All Negative (22/22) |
| **EV-B** | Coxsackievirus B1 (CV-B1), CV-B2, CV-B3, CV-B4, CV-B5, CV-B6, CV-A9, Echovirus 1 (E-1), E-2, E-3, E-4, E-5, E-6, E-7, E-9, E-11, E-12, E-13, E-14, E-15, E-16, E-17, E-18, E-19, E-20, E-21, E-24, E-25, E-26, E-27, E-29, E-30, E-31, E-32, E-33, enterovirus B69 (EV-B69), EV-B73, EV-B74, EV-B75, EV-B77, EV-B78, EV-B79, EV-B80, EV-B81, EV-B83, EV-B84, EV-B85, EV-B86, EV-B87, EV-B88, EV-B97, EV-B100, EV-B101, EV-B111, EV-B112, EV-B114 | All Negative (56/56) |
| **EV-C** | Poliovirus (PV) 1, PV-2, PV-3, (and Sabin PV 1, 2, 3), coxsackievirus A1 (CV-A1), CV-A11, CV-A13, CV-A17, CV-A19, CV-A20, CV-A21, CV-A22, CV-A24, EV-C96, EV-C99, EV-C102, EV-C105, EV-C113 | All Negative (21/21) |
| **EV-D (except EV-D68)** | EV-D70 (prototype J670/1971), EV-D94 (Nigeria, 2010 [two different isolates]), EV-D111 (Angola, 2012 [two different isolates]) | All Negative (5/5) |

[a] Three EV-A types, EV-A119, A120, and A121, are not available in CDC's collection or in public biorepositories. *In silico* examination of the missing EV-A nucleic acid sequences indicated that these viruses would not be detected by the pan-EV-D68 rRT-PCR assay.

[b] Seven EV-B types, EV-B82, B93, B98, B106, B107, B110, and B113, are not available in CDC's collection or in public biorepositories. *In silico* examination of the missing EV-B nucleic acid sequences indicated that these viruses would not be detected by the pan-EV-D68 rRT-PCR assay.

[c] Six EV-C types, EV-C95, C104, C109, C116, C117, and C118, are not available in CDC's collection or in public biorepositories. *In silico* examination of the missing EV-C nucleic acid sequences indicated that these viruses would not be detected by the pan-EV-D68 rRT-PCR assay.

[d] EV-D120, isolated from gorillas and a chimpanzee, in Cameroon and the Democratic Republic of Congo, respectively, was not available for testing. *In silico* examination of the EV-D120 nucleic acid sequences indicated that this virus would not be detected by the pan-EV-D68 rRT-PCR assay. To date, there are no reports of EV-D120 detection in humans.

**Rhinovirus Species A and B.** Rhinoviruses (RV) (genus *Enterovirus*) circulate widely during EV season and are commonly present in respiratory specimens. Therefore, we tested available RVs, using undiluted RNA extracted from frozen cell culture supernatants. A total of 96 culture-propagated RV, including 71 species A (RV-A) and 25 species B (RV-B) were tested (**Table S2**). The Pan-EV-D68 rRT-PCR did not amplify any viruses from RV-A or RV-B.

**Table S5: Pan-EV-D68 rRT-PCR analytical specificity confirmed using undiluted RNA extracted from rhinovirus (RV) cell culture supernatants**

| Species | Rhinovirus Types | Pan-EV-D68 rRT-PCR Result |
|---|---|---|
| **RV-A** | RV-A1, A2, A7, A8, A9, A10, A11, A12, A13, A15, A16, A18, A19, A20, A21, A22, A23, A24, A25, A28, A29, A30, A31, A32, A33, A34, A36, A38, A39, A40, A41, A43, A45, A46, A47, A49, A50, A51, A53, A54, A55, A56, A57, A58, A59, A60, A61, A62, A63, A64, A65, A66, A67, A68, A71, A73, A74, A75, A76, A77, A78, A80, A81, A82, A85, A88, A89, A90, A94, A96, A100 | All Negative (71/71) |
| **RV-B** | RV-B3, B4, B5, B6, B14, B17, B26, B27, B35, B37, B42, B48, B52, B69, B70, B72, B79, B83, B84, B86, B91, B92, B93, B97, B99 | All Negative (25/25) |

[a] Nine RV-A types, RV-A101, A102, A103, A104, A105, A106, A107, A108, and A109, are not available in CDC's collection or in public biorepositories. *In silico* examination of the missing RV-A nucleic acid sequences indicated that these viruses would not be detected by the pan-EV-D68 rRT-PCR assay.

[b] Seven RV-B types, RV-B100, B101, B102, B103, B104, B105, and B106, are not available in CDC's collection or in public biorepositories. *In silico* examination of the missing RV-B nucleic acid sequences indicated that these viruses would not be detected by the pan-EV-D68 rRT-PCR assay.

**Rhinovirus Species C.** Rhinovirus species C (RV-C) (genus *Enterovirus*) have not been successfully grown in conventional cell cultures. Sixty-nine RV-C, identified by direct sequencing of clinical specimens during 2014, were used to test specificity. Extracted RNA from respiratory tract clinical specimens (nasopharyngeal/oropharyngeal swabs in viral transport medium), containing RV-C types, was used to test the pan-EV-D68 rRT-PCR assay specificity. RV-C types in the NP/OP specimens were determined by Sanger sequencing of the VP1 region or by sequencing the RV VP4/VP2 region. The Pan-EV-D68 rRT-PCR did not amplify any of the RV-C viruses tested (**Table S3**).

**Table S6: Pan-EV-D68 rRT-PCR analytical specificity confirmed using RV-C clinical specimen RNA**

| *Enterovirus* Species | Rhinovirus Types | Pan-EV-D68 rRT-PCR Result |
|---|---|---|
| **RV-C** | RV-C2, C3, C4, C5, C6, C7, C9, C10, C11, C12, C14, C16, C18, C19, C21, C22, C23, C24, C25, C26, C27, C28, C29, C30, C31, C32, C33, C34, C35, C37, C38, C39, C40, C41, C42, C43, C44, C45 | All Negative (69/69) |

[a] Eighteen RV-C types, RV-C1, C8, C13, C15, C17, C20, C36, C46, C47, C48, C49, C50, C51, C52, C53, C54, C55 and C56, are not available in CDC's collection or in public biorepositories. -*In silico* examination of the missing RV-C nucleic acid sequences indicated that these viruses would not be detected by the pan-EV-D68 rRT-PCR assay.

**Other common respiratory viruses.** Extracted RNA from fourteen common respiratory viruses was tested using the pan-EV-D68 rRT-PCR. RNA was extracted from viral cell culture supernatants and tested undiluted. The Pan-EV-D68 rRT-PCR did not amplify any of the tested viruses (**Table S4**).

**Table S7: Pan-EV-D68 rRT-PCR cross-reactivity with common respiratory viruses**

| Virus[a] | Pan-EV-D68 rRT-PCR Result |
|---|---|
| **Adenovirus C1 (AdV-71)** | Negative |
| **CoV 229E** | Negative |
| **CoV OC43** | Negative |
| **CoV MERS** | Negative |
| **SARS-CoV-2** | Negative |
| **HMPV (CAN99-81)** | Negative |
| **Influenza A H1N1** | Negative |
| **Influenza A H3N2** | Negative |
| **Influenza B (B/Vic)** | Negative |
| **PIV1 (C35)** | Negative |
| **PIV2 (Greer)** | Negative |
| **PIV3 (C-43)** | Negative |
| **PIV4a (CH 19503)** | Negative |
| **RSV-A (Long)** | Negative |

[a] CoV, coronavirus; HMPV, human metapneumovirus; PIV, parainfluenza virus; RSV, respiratory syncytial virus.

**Other organisms commonly detected in respiratory tract specimens.** Additional evaluation of the analytical specificity of the pan-EV-D68 rRT-PCR was performed through *in silico* analysis of the CDC2015 primer and probe sequences against common causes of respiratory illness and other clinically relevant organisms. BLASTn analysis queries of the VP1.2020 primers and probe were performed against the GenBank public domain nucleotide sequences and showed no significant combined homologies (primer targets and probe target) with other conditions that would predict potential false positive rRT-PCR results.

**Table S8: CDC2015 primer and probe set *in silico* specificity analysis**

| Disease/condition | Taxid | Agent |
|---|---|---|
| **Human parainfluenza** | 11226 | Parainfluenza virus 4b |
| **Human parechovirus** | 12063 | Parechovirus A1 |
| **Common cold, bronchiolitis, pneumonia** | 208895 | Respiratory syncytial virus B |
| **Sinusitis** | 1280 | *Staphylococcus aureus* |
| **Pertussis (whooping cough)** | 520 | *Bordetella pertussis* |
| **Mycoplasma pneumonia** | 2104 | *Mycoplasma pneumoniae* |
| **Pharyngitis, bronchitis, atypical pneumonia** | 83558 | *Chlamydia pneumoniae*<br>(or *Chlamydophila pneumonia*) |
| **Bacteremia, pneumonia, epiglottitis and acute bacterial meningitis** | 727 | *Haemophilus influenza* |
| **Tuberculosis** | 1773 | *Mycobacterium tuberculosis* |
| **Lyme disease** | 139 | *Borrelia burgdorferi* |
| **Respiratory infection** | 550 | *Enterobacter cloacae* |
| **Septicaemia** | 1354 | *Enterococcus hirae* |
| **Foodborne illness** | 1502 | *Clostridium perfringens* |
| **Abdominal infection (rare)** | 46503 | *Parabacteroides merdae* |
| **Human gastrointestinal bacterium** | 1680 | *Bifidobacterium adolescentis* |
| **Human gastrointestinal bacterium** | 216816 | *Bifidobacterium longum* |
| **Human gastrointestinal bacterium** | 40518 | *Ruminococcus bromii* |

**Table S9. Line-by-line description of respiratory clinical samples tested by both snPCR/Seq and CDC2022 rRT-PCR.**

| Number | VP1 RT-snPCR/Seq Result | Pan-EV-D68 CDC2022 Result (Ct or positivity) | Year |
|---|---|---|---|
| 1 | E-11 | Negative | 2015 |
| 2 | CV-A6 | Negative | 2015 |
| 3 | RV-80 | Negative | 2015 |
| 4 | CV-A6 | Negative | 2015 |
| 5 | CV-A4 | Negative | 2015 |
| 6 | E-11 | Negative | 2015 |
| 7 | CV-A6 | Negative | 2015 |
| 8 | E-3 | Negative | 2015 |
| 9 | E-25 | Negative | 2016 |
| 10 | Negative | Negative | 2016 |
| 11 | RV-A57 | Negative | 2016 |
| 12 | Negative | Negative | 2016 |
| 13 | CV-A6 | Negative | 2016 |
| 14 | RV-A51 | Negative | 2016 |
| 15 | RV-A21 | Negative | 2016 |
| 16 | CV-A6 | Negative | 2016 |
| 17 | Negative | Negative | 2016 |
| 18 | Negative | Negative | 2016 |
| 19 | CV-A9 | Negative | 2016 |
| 20 | CV-A9 | Negative | 2016 |
| 21 | E-6 | Negative | 2016 |
| 22 | Negative | Negative | 2016 |
| 23 | CV-A6 | Negative | 2016 |
| 24 | Negative | Negative | 2016 |
| 25 | RV-A49 | Negative | 2016 |
| 26 | Negative | Negative | 2016 |
| 27 | Negative | Negative | 2016 |
| 28 | CV-A6 | Negative | 2016 |
| 29 | RV-A20 | Negative | 2016 |
| 30 | RV-A12 | Negative | 2016 |
| 31 | E-6 | Negative | 2016 |
| 32 | Negative | Negative | 2016 |
| 33 | Negative | Negative | 2017 |
| 34 | Negative | Negative | 2017 |
| 35 | RV-A49 | Negative | 2017 |
| 36 | CV-B3 | Negative | 2017 |
| 37 | Negative | Negative | 2017 |
| 38 | Negative | Negative | 2017 |
| 39 | Negative | Negative | 2017 |
| 40 | Negative | Negative | 2017 |
| 41 | RV-A101 | Negative | 2017 |
| 42 | RV-A56 | Negative | 2017 |
| 43 | RV-A55 | Negative | 2017 |
| 44 | CV-A6 | Negative | 2017 |
| 45 | CV-B2 | Negative | 2017 |
| 46 | RV-A1B | Negative | 2017 |

| | | | |
|---|---|---|---|
| 47 | Negative | Negative | 2017 |
| 48 | CV-B5 | Negative | 2017 |
| 49 | RV-A24 | Negative | 2017 |
| 50 | RV-A45 | Negative | 2018 |
| 51 | EV-D68 | 19.9 | 2014 |
| 52 | EV-D68 | 26.1 | 2014 |
| 53 | Negative | Negative | 2014 |
| 54 | EV-D68 | 29.8 | 2014 |
| 55 | EV-D68 | 17.6 | 2014 |
| 56 | Negative | Negative | 2014 |
| 57 | Negative | Negative | 2014 |
| 58 | EV-D68 | 18.7 | 2014 |
| 59 | Negative | Negative | 2014 |
| 60 | Negative | Negative | 2014 |
| 61 | EV-D68 | 27.6 | 2014 |
| 62 | EV-D68 | 29.5 | 2014 |
| 63 | EV-D68 | 29.2 | 2014 |
| 64 | Negative | Negative | 2014 |
| 65 | EV-D68 | 26.7 | 2014 |
| 66 | EV-D68 | 16.8 | 2014 |
| 67 | EV-D68 | 23.2 | 2014 |
| 68 | EV-D68 | 29.9 | 2014 |
| 69 | EV-D68 | 21.8 | 2014 |
| 70 | RV-B27 | Negative | 2014 |
| 71 | EV-D68 | 20.7 | 2014 |
| 72 | EV-D68 | 27.9 | 2014 |
| 73 | EV-D68 | 19.6 | 2014 |
| 74 | EV-D68 | 32.3 | 2014 |
| 75 | Negative | Negative | 2014 |
| 76 | EV-D68 | 31.4 | 2014 |
| 77 | Negative | Negative | 2014 |
| 78 | EV-D68 | 22.5 | 2014 |
| 79 | EV-D68 | 23 | 2014 |
| 80 | Negative | Negative | 2014 |
| 81 | EV-D68 | 27.8 | 2014 |
| 82 | EV-D68 | 19.1 | 2014 |
| 83 | EV-D68 | 23.5 | 2014 |
| 84 | RV-A101 | Negative | 2014 |
| 85 | RV-A101 | Negative | 2014 |
| 86 | CV-A6 | Negative | 2014 |
| 87 | Negative | Negative | 2014 |
| 88 | EV-D68 | 15.9 | 2014 |
| 89 | CV-A6 | Negative | 2014 |
| 90 | RV-77 | Negative | 2014 |
| 91 | EV-A71 | Negative | 2014 |
| 92 | EV-D68 | 20.2 | 2014 |
| 93 | EV-D68 | 15.6 | 2014 |
| 94 | EV-D68 | 24.9 | 2014 |
| 95 | EV-D68 | 20.1 | 2014 |
| 96 | EV-D68 | 20.2 | 2014 |
| 97 | EV-D68 | 20.4 | 2014 |

| | | | |
|---|---|---|---|
| 98 | EV-D68 | 30.6 | 2014 |
| 99 | EV-D68 | 26.2 | 2014 |
| 100 | RV-59 | Negative | 2014 |
| 101 | RV-A94 | Negative | 2014 |
| 102 | EV-D68 | 29.1 | 2014 |
| 103 | EV-D68 | 31.9 | 2014 |
| 104 | EV-D68 | 28.9 | 2014 |
| 105 | EV-D68 | 26.4 | 2014 |
| 106 | EV-D68 | 21.8 | 2014 |
| 107 | EV-D68 | 28.3 | 2014 |
| 108 | EV-D68 | 25 | 2014 |
| 109 | EV-D68 | 26.8 | 2014 |
| 110 | EV-D68 | 35.6 | 2014 |
| 111 | RV-A34 | Negative | 2014 |
| 112 | RV-A63 | Negative | 2014 |
| 113 | EV-D68 | 29.9 | 2014 |
| 114 | RV-A49 | Negative | 2014 |
| 115 | E-9 | Negative | 2014 |
| 116 | EV-D68 | 21 | 2014 |
| 117 | EV-A71 | Negative | 2014 |
| 118 | EV-D68 | 25.1 | 2014 |
| 119 | RV-A24 | Negative | 2014 |
| 120 | EV-D68 | 18.2 | 2014 |
| 121 | EV-D68 | 20.9 | 2014 |
| 122 | EV-D68 | 18.4 | 2014 |
| 123 | EV-D68 | 15.6 | 2014 |
| 124 | EV-D68 | 28.9 | 2014 |
| 125 | EV-D68 | 23.8 | 2014 |
| 126 | RV-C | Negative | 2014 |
| 127 | RV-C | Negative | 2014 |
| 128 | EV-D68 | 28.6 | 2014 |
| 129 | RV-B6 | Negative | 2014 |
| 130 | RV-B6 | Negative | 2014 |
| 131 | RV-A101 | Negative | 2014 |
| 132 | EV-D68 | 20.5 | 2014 |
| 133 | EV-D68 | 24.3 | 2014 |
| 134 | EV-D68 | 27.5 | 2014 |
| 135 | EV-D68 | 22 | 2014 |
| 136 | EV-D68 | 23.9 | 2014 |
| 137 | RV-A73 | Negative | 2014 |
| 138 | Negative | Negative | 2014 |
| 139 | EV-D68 | 29.2 | 2014 |
| 140 | RV-B48 | Negative | 2014 |
| 141 | CV-A10 | Negative | 2014 |
| 142 | EV-D68 | 27.3 | 2014 |
| 143 | EV-D68 | 21.4 | 2014 |
| 144 | EV-D68 | 28.9 | 2014 |
| 145 | EV-D68 | 26.2 | 2014 |
| 146 | EV-D68 | 26.6 | 2014 |
| 147 | EV-D68 | 29.8 | 2014 |
| 148 | EV-D68 | 17.4 | 2014 |

| | | | |
|---|---|---|---|
| 149 | EV-D68 | 33.5 | 2014 |
| 150 | EV-D68 | 32.5 | 2014 |
| 151 | EV-D68 | 31.1 | 2014 |
| 152 | EV-D68 | 25.3 | 2014 |
| 153 | EV-D68 | 24.8 | 2014 |
| 154 | EV-D68 | 19.9 | 2014 |
| 155 | EV-D68 | 17.2 | 2014 |
| 156 | Negative | Negative | 2014 |
| 157 | Negative | Negative | 2014 |
| 158 | EV-D68 | 21.5 | 2014 |
| 159 | Negative | Negative | 2014 |
| 160 | EV-D68 | 15.7 | 2014 |
| 161 | Negative | Negative | 2014 |
| 162 | EV-D68 | 31.3 | 2014 |
| 163 | EV-D68 | 23 | 2014 |
| 164 | EV-D68 | 20.3 | 2014 |
| 165 | EV-D68 | 26.3 | 2014 |
| 166 | EV-D68 | 30.5 | 2014 |
| 167 | EV-D68 | 33.3 | 2014 |
| 168 | EV-D68 | 34 | 2014 |
| 169 | EV-D68 | 21.6 | 2014 |
| 170 | EV-D68 | 24.5 | 2014 |
| 171 | RV-B37 | 31.5 | 2014 |
| 172 | Negative | Negative | 2014 |
| 173 | Negative | Negative | 2014 |
| 174 | Negative | Negative | 2014 |
| 175 | EV-D68 | 31.1 | 2014 |
| 176 | EV-D68 | 29.2 | 2016 |
| 177 | EV-D68 | 30.3 | 2016 |
| 178 | EV-D68 | 33.9 | 2016 |
| 179 | EV-D68 | 32.7 | 2016 |
| 180 | EV-D68 | 32.6 | 2016 |
| 181 | EV-D68 | 38.9 | 2016 |
| 182 | EV-D68 | 32 | 2016 |
| 183 | EV-D68 | 30 | 2016 |
| 184 | EV-D68 | 35.5 | 2016 |
| 185 | EV-D68 | 29.4 | 2016 |
| 186 | EV-D68 | 26.6 | 2016 |
| 187 | EV-D68 | 27 | 2016 |
| 188 | EV-D68 | 21.6 | 2016 |
| 189 | EV-D68 | 28.4 | 2016 |
| 190 | EV-D68 | 25.4 | 2016 |
| 191 | EV-D68 | 22 | 2016 |
| 192 | EV-D68 | 15 | 2016 |
| 193 | EV-D68 | 27.9 | 2016 |
| 194 | EV-D68 | 20.1 | 2016 |
| 195 | EV-D68 | 21 | 2016 |
| 196 | EV-D68 | 20.3 | 2016 |
| 197 | EV-D68 | 32.2 | 2016 |
| 198 | EV-D68 | 25.4 | 2016 |
| 199 | EV-D68 | 28.1 | 2016 |

| | | | |
|---|---|---|---|
| 200 | EV-D68 | 23.6 | 2016 |
| 201 | EV-D68 | 18.7 | 2016 |
| 202 | EV-D68 | 22.7 | 2016 |
| 203 | EV-D68 | 30.6 | 2016 |
| 204 | EV-D68 | 25.2 | 2016 |
| 205 | EV-D68 | 28.9 | 2016 |
| 206 | EV-D68 | 27.8 | 2016 |
| 207 | EV-D68 | 27.8 | 2016 |
| 208 | EV-D68 | 26.7 | 2016 |
| 209 | EV-D68 | 33 | 2016 |
| 210 | EV-D68 | 26.9 | 2016 |
| 211 | EV-D68 | 30.8 | 2016 |
| 212 | EV-D68 | 22.2 | 2016 |
| 213 | EV-D68 | 27.8 | 2016 |
| 214 | EV-D68 | 31.4 | 2016 |
| 215 | EV-D68 | 18.6 | 2016 |
| 216 | EV-D68 | 27.4 | 2016 |
| 217 | EV-D68 | 30.4 | 2016 |
| 218 | EV-D68 | 29.6 | 2016 |
| 219 | EV-D68 | 32.7 | 2016 |
| 220 | EV-D68 | 19.7 | 2016 |
| 221 | EV-D68 | 29 | 2016 |
| 222 | EV-D68 | 32 | 2016 |
| 223 | EV-D68 | 28.2 | 2016 |
| 224 | EV-D68 | 26 | 2016 |
| 225 | EV-D68 | 34.2 | 2016 |
| 226 | EV-D68 | 27.3 | 2016 |
| 227 | EV-D68 | 35 | 2016 |
| 228 | EV-D68 | 24.1 | 2016 |
| 229 | EV-D68 | 30.8 | 2016 |
| 230 | EV-D68 | 30.1 | 2016 |
| 231 | EV-D68 | 29.6 | 2016 |
| 232 | EV-D68 | 23.6 | 2016 |
| 233 | EV-D68 | 28.6 | 2016 |
| 234 | EV-D68 | 22.5 | 2016 |
| 235 | EV-D68 | 21.7 | 2016 |
| 236 | EV-D68 | 24.4 | 2016 |
| 237 | EV-D68 | 31.4 | 2016 |
| 238 | EV-D68 | 30.8 | 2016 |
| 239 | EV-D68 | 37.6 | 2016 |
| 240 | EV-D68 | 29.4 | 2016 |
| 241 | EV-D68 | 29.2 | 2016 |
| 242 | EV-D68 | 28.8 | 2016 |
| 243 | EV-D68 | 29.6 | 2016 |
| 244 | EV-D68 | 29.9 | 2016 |
| 245 | EV-D68 | 31.6 | 2016 |
| 246 | EV-D68 | 36.5 | 2016 |
| 247 | EV-D68 | 31.3 | 2016 |
| 248 | EV-D68 | 37.8 | 2016 |
| 249 | EV-D68 | 28.6 | 2016 |
| 250 | EV-D68 | 26.7 | 2016 |

| | | | |
|---|---|---|---|
| 251 | EV-D68 | 26.6 | 2016 |
| 252 | EV-D68 | 23.8 | 2016 |
| 253 | EV-D68 | 22 | 2017 |
| 254 | EV-D68 | 26 | 2017 |
| 255 | EV-D68 | 26.8 | 2017 |
| 256 | EV-D68 | 25.6 | 2017 |
| 257 | EV-D68 | 22.8 | 2017 |
| 258 | EV-D68 | 22.4 | 2017 |
| 259 | EV-D68 | 26.6 | 2017 |
| 260 | EV-D68 | 27 | 2017 |
| 261 | EV-D68 | 28.5 | 2017 |
| 262 | EV-D68 | 23 | 2017 |
| 263 | EV-D68 | 28.9 | 2017 |
| 264 | EV-D68 | 32.6 | 2018 |
| 265 | EV-D68 | 32 | 2018 |
| 266 | EV-D68 | 23.5 | 2018 |
| 267 | EV-D68 | 26.9 | 2018 |
| 268 | EV-D68 | 21 | 2012 |
| 269 | RV-A45 | Negative | 2018 |
| 270 | Negative | Negative | 2018 |
| 271 | Negative | Negative | 2018 |
| 272 | Negative | Negative | 2018 |
| 273 | Negative | Negative | 2018 |
| 274 | Negative | Negative | 2018 |
| 275 | EV-D68 | 36.02 | 2018 |
| 276 | EV-D68 | 35.55 | 2018 |
| 277 | EV-D68 | 28.78 | 2018 |
| 278 | EV-D68 | 31 | 2018 |
| 279 | Negative | Negative | 2018 |
| 280 | Negative | Negative | 2018 |
| 281 | Negative | Negative | 2018 |
| 282 | EV-D68 | 25.03 | 2018 |
| 283 | Negative | Negative | 2018 |
| 284 | Negative | Negative | 2018 |
| 285 | Negative | Negative | 2018 |
| 286 | Negative | Negative | 2018 |
| 287 | Negative | Negative | 2018 |
| 288 | Negative | Negative | 2018 |
| 289 | EV-A71 | Negative | 2018 |
| 290 | EV-A71 | Negative | 2018 |
| 291 | EV-A71 | Negative | 2018 |
| 292 | Negative | Negative | 2018 |
| 293 | Negative | Negative | 2018 |
| 294 | Negative | Negative | 2018 |
| 295 | Negative | Negative | 2018 |
| 296 | Negative | Negative | 2018 |
| 297 | Negative | Negative | 2018 |
| 298 | Negative | Negative | 2018 |
| 299 | Negative | Negative | 2018 |
| 300 | Negative | Negative | 2018 |
| 301 | Negative | Negative | 2018 |

| | | | |
|---|---|---|---|
| 302 | Negative | Negative | 2018 |
| 303 | Negative | Negative | 2018 |
| 304 | RV-A81 | Negative | 2018 |
| 305 | EV-D68 | 26.54 | 2018 |
| 306 | EV-D68 | 31.95 | 2018 |
| 307 | Negative | Negative | 2018 |
| 308 | EV-D68 | 24.25 | 2018 |
| 309 | Negative | Negative | 2018 |
| 310 | EV-D68 | 23.24 | 2018 |
| 311 | Negative | Negative | 2018 |
| 312 | EV-D68 | 29.38 | 2018 |
| 313 | EV-D68 | 29.86 | 2018 |
| 314 | Negative | Negative | 2018 |
| 315 | EV-D68 | 28.9 | 2018 |
| 316 | Negative | Negative | 2018 |
| 317 | EV-D68 | 35.11 | 2018 |
| 318 | Negative | Negative | 2018 |
| 319 | Negative | Negative | 2018 |
| 320 | Negative | Negative | 2018 |
| 321 | RV-B4 | Negative | 2018 |
| 322 | Negative | Negative | 2018 |
| 323 | Negative | Negative | 2018 |
| 324 | Negative | Negative | 2018 |
| 325 | Negative | Negative | 2018 |
| 326 | Negative | Negative | 2018 |
| 327 | Negative | Negative | 2018 |
| 328 | RV-A56 | Negative | 2018 |
| 329 | EV-D68 | 29.62 | 2018 |
| 330 | RV-B104 | Negative | 2018 |
| 331 | EV-D68 | 18.27 | 2018 |
| 332 | RV-A38 | Negative | 2018 |
| 333 | EV-A71 | Negative | 2018 |
| 334 | CV-A5 | Negative | 2018 |
| 335 | Negative | Negative | 2018 |
| 336 | RV-A40 | Negative | 2018 |
| 337 | EV-A71 | Negative | 2018 |
| 338 | EV-A71 | Negative | 2018 |
| 339 | EV-D68 | 17.6 | 2018 |
| 340 | EV-D68 | 20.9 | 2018 |
| 341 | EV-D68 | 18.6 | 2018 |
| 342 | EV-D68 | 18.9 | 2018 |
| 343 | EV-D68 | 17.9 | 2018 |
| 344 | EV-D68 | 19.5 | 2018 |
| 345 | EV-D68 | 19.7 | 2018 |
| 346 | EV-D68 | 32.1 | 2018 |
| 347 | EV-D68 | 22.6 | 2018 |
| 348 | EV-D68 | 15.6 | 2018 |
| 349 | Negative | Negative | 2018 |
| 350 | EV-D68 | 27.28 | 2018 |
| 351 | RV-A90 | Negative | 2018 |
| 352 | EV-D68 | 31 | 2018 |

| | | | |
|---|---|---|---|
| 353 | Negative | Negative | 2018 |
| 354 | Negative | Negative | 2018 |
| 355 | EV-D68 | 36.61 | 2018 |
| 356 | EV-D68 | 30.82 | 2018 |
| 357 | Negative | Negative | 2018 |
| 358 | RV-A54 | Negative | 2018 |
| 359 | Negative | Negative | 2018 |
| 360 | RV-A101 | Negative | 2018 |
| 361 | RV-B4 | Negative | 2018 |
| 362 | Negative | Negative | 2018 |
| 363 | Negative | Negative | 2018 |
| 364 | EV-D68 | 25.14 | 2018 |
| 365 | Negative | Negative | 2018 |
| 366 | EV-D68 | 31.84 | 2018 |
| 367 | EV-D68 | 30.57 | 2018 |
| 368 | Negative | Negative | 2018 |
| 369 | Negative | Negative | 2018 |
| 370 | EV-D68 | 30.96 | 2018 |
| 371 | Negative | Negative | 2018 |
| 372 | Negative | Negative | 2018 |
| 373 | Negative | Negative | 2018 |
| 374 | Negative | Negative | 2018 |
| 375 | EV-D68 | 24.04 | 2018 |
| 376 | EV-D68 | 31.94 | 2018 |
| 377 | Negative | Negative | 2018 |
| 378 | Negative | Negative | 2018 |
| 379 | Negative | Negative | 2018 |
| 380 | Negative | Negative | 2018 |
| 381 | Negative | Negative | 2018 |
| 382 | Negative | Negative | 2018 |
| 383 | Negative | Negative | 2018 |
| 384 | Negative | Negative | 2018 |
| 385 | Negative | Negative | 2018 |
| 386 | Negative | Negative | 2018 |
| 387 | EV-D68 | 32.11 | 2018 |
| 388 | Negative | Negative | 2018 |
| 389 | Negative | Negative | 2018 |
| 390 | Negative | Negative | 2018 |
| 391 | EV-D68 | 29.18 | 2018 |
| 392 | EV-D68 | 36.23 | 2018 |
| 393 | Negative | Negative | 2018 |
| 394 | Negative | Negative | 2018 |
| 395 | Negative | Negative | 2018 |
| 396 | Negative | Negative | 2018 |
| 397 | EV-D68 | 24.63 | 2018 |
| 398 | Negative | Negative | 2018 |
| 399 | EV-D68 | 35.46 | 2018 |
| 400 | Negative | Negative | 2018 |
| 401 | Negative | Negative | 2018 |
| 402 | RV-B4 | Negative | 2018 |
| 403 | Negative | Negative | 2018 |

| | | | |
|---|---|---|---|
| 404 | Negative | Negative | 2018 |
| 405 | Negative | Negative | 2018 |
| 406 | Negative | Negative | 2018 |
| 407 | EV-D68 | 26.46 | 2018 |
| 408 | Negative | Negative | 2018 |
| 409 | Negative | Negative | 2018 |
| 410 | Negative | Negative | 2018 |
| 411 | Negative | Negative | 2018 |
| 412 | Negative | Negative | 2018 |
| 413 | Negative | Negative | 2018 |
| 414 | EV-D68 | 24.95 | 2018 |
| 415 | Negative | Negative | 2018 |
| 416 | CV-A6 | Negative | 2018 |
| 417 | CV-A6 | Negative | 2018 |
| 418 | Negative | Negative | 2018 |
| 419 | Negative | Negative | 2018 |
| 420 | CV-A6 | Negative | 2018 |
| 421 | Negative | Negative | 2018 |
| 422 | Negative | Negative | 2018 |
| 423 | Negative | Negative | 2018 |
| 424 | RV-A85 | Negative | 2018 |
| 425 | Negative | Negative | 2018 |
| 426 | Negative | Negative | 2018 |
| 427 | RV-A104 | Negative | 2018 |
| 428 | Negative | Negative | 2018 |
| 429 | Negative | Negative | 2018 |
| 430 | Negative | Negative | 2018 |
| 431 | Negative | Negative | 2018 |
| 432 | Negative | Negative | 2018 |
| 433 | Negative | Negative | 2018 |
| 434 | Negative | Negative | 2018 |
| 435 | Negative | Negative | 2018 |
| 436 | EV-D68 | 32.7 | 2018 |
| 437 | Negative | Negative | 2018 |
| 438 | Negative | Negative | 2018 |
| 439 | Negative | Negative | 2018 |
| 440 | Negative | Negative | 2018 |
| 441 | Negative | Negative | 2018 |
| 442 | Negative | Negative | 2018 |
| 443 | Negative | Negative | 2018 |
| 444 | Negative | Negative | 2018 |
| 445 | Negative | Negative | 2018 |
| 446 | EV-D68 | 29.7 | 2018 |
| 447 | Negative | Negative | 2018 |
| 448 | Negative | Negative | 2018 |
| 449 | Negative | Negative | 2018 |
| 450 | Negative | Negative | 2018 |
| 451 | Negative | Negative | 2018 |
| 452 | Negative | Negative | 2018 |
| 453 | Negative | Negative | 2018 |
| 454 | Negative | Negative | 2018 |

| | | | |
|---|---|---|---|
| 455 | Negative | Negative | 2018 |
| 456 | EV-D68 | 25.1 | 2018 |
| 457 | RV-A101 | Negative | 2018 |
| 458 | RV-A22 | Negative | 2018 |
| 459 | Negative | Negative | 2018 |
| 460 | EV-D68 | 28.9 | 2018 |
| 461 | EV-D68 | 33.6 | 2018 |
| 462 | RV-A12 | Negative | 2018 |
| 463 | Negative | Negative | 2018 |
| 464 | EV-D68 | 29.3 | 2018 |
| 465 | EV-D68 | 32.9 | 2018 |
| 466 | Negative | Negative | 2018 |
| 467 | Negative | Negative | 2018 |
| 468 | Negative | Negative | 2018 |
| 469 | Negative | Negative | 2018 |
| 470 | Negative | Negative | 2018 |
| 471 | Negative | Negative | 2018 |
| 472 | Negative | Negative | 2018 |
| 473 | EV-D68 | 32.6 | 2018 |
| 474 | Negative | Negative | 2018 |
| 475 | Negative | Negative | 2018 |
| 476 | EV-D68 | 30 | 2018 |
| 477 | EV-D68 | 27.5 | 2018 |
| 478 | Negative | Negative | 2018 |
| 479 | Negative | Negative | 2018 |
| 480 | Negative | Negative | 2018 |
| 481 | E-11 | Negative | 2018 |
| 482 | EV-A71 | Negative | 2018 |
| 483 | Negative | Negative | 2018 |
| 484 | Negative | Negative | 2018 |
| 485 | Negative | Negative | 2018 |
| 486 | EV-D68 | 33.3 | 2018 |
| 487 | Negative | Negative | 2018 |
| 488 | Negative | Negative | 2018 |
| 489 | Negative | Negative | 2018 |
| 490 | Negative | Negative | 2018 |
| 491 | Negative | Negative | 2018 |
| 492 | Negative | Negative | 2018 |
| 493 | Negative | Negative | 2018 |
| 494 | Negative | Negative | 2018 |
| 495 | CV-B2 | Negative | 2018 |
| 496 | Negative | Negative | 2018 |
| 497 | Negative | Negative | 2018 |
| 498 | RV-A8 | Negative | 2018 |
| 499 | EV-D68 | 30.9 | 2018 |
| 500 | Negative | Negative | 2018 |
| 501 | Negative | Negative | 2018 |
| 502 | Negative | Negative | 2018 |
| 503 | Negative | Negative | 2018 |
| 504 | Negative | Negative | 2018 |
| 505 | Negative | Negative | 2018 |

| | | | |
|---|---|---|---|
| 506 | Negative | Negative | 2018 |
| 507 | RV-A16 | Negative | 2018 |
| 508 | Negative | Negative | 2018 |
| 509 | Negative | Negative | 2018 |
| 510 | Negative | Negative | 2018 |
| 511 | Negative | Negative | 2018 |
| 512 | Negative | Negative | 2018 |
| 513 | Negative | Negative | 2018 |
| 514 | Negative | Negative | 2018 |
| 515 | Negative | Negative | 2018 |
| 516 | Negative | Negative | 2018 |
| 517 | Negative | Negative | 2018 |
| 518 | CV-A6 | Negative | 2018 |
| 519 | Negative | Negative | 2018 |
| 520 | Negative | Negative | 2019 |
| 521 | Negative | Negative | 2019 |
| 522 | Negative | Negative | 2019 |
| 523 | Negative | Negative | 2019 |
| 524 | Negative | Negative | 2019 |
| 525 | Negative | Negative | 2019 |
| 526 | Negative | Negative | 2019 |
| 527 | Negative | Negative | 2019 |
| 528 | Negative | Negative | 2019 |
| 529 | Negative | Negative | 2019 |
| 530 | Negative | Negative | 2019 |
| 531 | Negative | Negative | 2019 |
| 532 | Negative | Negative | 2019 |
| 533 | Negative | Negative | 2019 |
| 534 | Negative | Negative | 2019 |
| 535 | Negative | Negative | 2019 |
| 536 | Negative | Negative | 2019 |
| 537 | Negative | Negative | 2019 |
| 538 | Negative | Negative | 2019 |
| 539 | Negative | Negative | 2019 |
| 540 | Negative | Negative | 2019 |
| 541 | Negative | Negative | 2019 |
| 542 | Negative | Negative | 2019 |
| 543 | Negative | Negative | 2019 |
| 544 | Negative | Negative | 2019 |
| 545 | Negative | Negative | 2019 |
| 546 | Negative | Negative | 2019 |
| 547 | Negative | Negative | 2019 |
| 548 | Negative | Negative | 2019 |
| 549 | Negative | Negative | 2019 |
| 550 | Negative | Negative | 2019 |
| 551 | Negative | Negative | 2019 |
| 552 | Negative | Negative | 2019 |
| 553 | Negative | Negative | 2019 |
| 554 | Negative | Negative | 2019 |
| 555 | Negative | Negative | 2019 |
| 556 | Negative | Negative | 2019 |

| | | | |
|---|---|---|---|
| 557 | Negative | Negative | 2019 |
| 558 | RV-A12 | Negative | 2019 |
| 559 | RV-A12 | Negative | 2019 |
| 560 | Negative | Negative | 2019 |
| 561 | Negative | Negative | 2019 |
| 562 | Negative | Negative | 2019 |
| 563 | Negative | Negative | 2019 |
| 564 | Negative | Negative | 2019 |
| 565 | Negative | Negative | 2019 |
| 566 | Negative | Negative | 2019 |
| 567 | Negative | Negative | 2019 |
| 568 | RV-A34 | Negative | 2019 |
| 569 | Negative | Negative | 2019 |
| 570 | Negative | Negative | 2019 |
| 571 | Negative | Negative | 2019 |
| 572 | Negative | Negative | 2019 |
| 573 | Negative | Negative | 2019 |
| 574 | Negative | Negative | 2019 |
| 575 | EV-D68 | EV-D68(+) | 2018 |
| 576 | EV-D68 | EV-D68(+) | 2018 |
| 577 | EV-D68 | EV-D68(+) | 2018 |
| 578 | EV-D68 | EV-D68(+) | 2018 |
| 579 | EV-D68 | EV-D68(+) | 2018 |
| 580 | EV-D68 | EV-D68(+) | 2018 |
| 581 | EV-D68 | EV-D68(+) | 2018 |
| 582 | EV-D68 | EV-D68(+) | 2018 |
| 583 | EV-D68 | EV-D68(+) | 2018 |
| 584 | EV-D68 | EV-D68(+) | 2018 |
| 585 | EV-D68 | EV-D68(+) | 2018 |
| 586 | EV-D68 | EV-D68(+) | 2018 |
| 587 | EV-D68 | EV-D68(+) | 2018 |
| 588 | EV-D68 | EV-D68(+) | 2018 |
| 589 | EV-D68 | EV-D68(+) | 2018 |
| 590 | EV-D68 | EV-D68(+) | 2018 |
| 591 | EV-D68 | EV-D68(+) | 2018 |
| 592 | EV-D68 | EV-D68(+) | 2018 |
| 593 | EV-D68 | EV-D68(+) | 2018 |
| 594 | EV-D68 | EV-D68(+) | 2018 |
| 595 | EV-D68 | EV-D68(+) | 2018 |
| 596 | EV-D68 | EV-D68(+) | 2018 |
| 597 | EV-D68 | EV-D68(+) | 2018 |
| 598 | EV-D68 | EV-D68(+) | 2018 |
| 599 | EV-D68 | EV-D68(+) | 2018 |
| 600 | EV-D68 | EV-D68(+) | 2018 |
| 601 | EV-D68 | EV-D68(+) | 2018 |
| 602 | EV-D68 | EV-D68(+) | 2018 |
| 603 | EV-D68 | EV-D68(+) | 2018 |
| 604 | EV-D68 | EV-D68(+) | 2018 |
| 605 | EV-D68 | EV-D68(+) | 2018 |
| 606 | EV-D68 | EV-D68(+) | 2018 |
| 607 | EV-D68 | EV-D68(+) | 2018 |

| | | | |
|---|---|---|---|
| 608 | EV-D68 | EV-D68(+) | 2018 |
| 609 | EV-D68 | EV-D68(+) | 2018 |
| 610 | EV-D68 | EV-D68(+) | 2018 |
| 611 | EV-D68 | EV-D68(+) | 2018 |
| 612 | EV-D68 | EV-D68(+) | 2018 |
| 613 | EV-D68 | EV-D68(+) | 2018 |
| 614 | EV-D68 | EV-D68(+) | 2019 |
| 615 | EV-D68 | EV-D68(+) | 2022 |
| 616 | EV-D68 | EV-D68(+) | 2022 |
| 617 | EV-D68 | EV-D68(+) | 2022 |
| 618 | EV-D68 | EV-D68(+) | 2022 |
| 619 | EV-D68 | EV-D68(+) | 2022 |
| 620 | EV-D68 | EV-D68(+) | 2022 |
| 621 | EV-D68 | EV-D68(+) | 2022 |
| 622 | EV-D68 | EV-D68(+) | 2022 |
| 623 | EV-D68 | EV-D68(+) | 2022 |
| 624 | EV-D68 | EV-D68(+) | 2022 |
| 625 | EV-D68 | EV-D68(+) | 2022 |
